# pLM-Guided Inverse Folding for Antibody Sequence Design

**DOI:** 10.64898/2026.06.04.730089

**Authors:** Valentin Noske, Félix Koulischer, Kathleen Marchal, Thomas Demeester

## Abstract

Inverse folding, predicting amino acid sequences from three-dimensional structures, is a foundational task in computational protein design, yet it is hindered by the scarcity of structural data, which limits model training and risks overfitting. The standard approach fine-tunes general inverse folding models on domain-specific structural datasets like antibodies, but such data remain expensive. To enable inverse folders to benefit more from abundant sequence data, we propose combining ProteinMPNN, a general protein inverse folding model, with IgLM, an antibody-specific language model, via a training-free weighted ensemble of their predictions at inference time. Evaluated on antibody and nanobody structures, our results show that this approach substantially improves amino acid recovery over ProteinMPNN alone, approaching the performance of antibody-specific models like AntiFold while generating more diverse sequences. Even models already fine-tuned on antibody structures (AbMPNN) benefit from language model guidance, demonstrating that it complements structural fine-tuning and leads to more natural-looking sequences that still satisfy structural constraints.

## Introduction

Inverse folding, the task of predicting an amino acid sequence conditioned on a three-dimensional backbone, remains a foundational component of modern *in silico* protein design pipelines (Watson et al., 2023; Frank et al., 2024; Pacesa et al., 2025). In these workflows, a backbone is generated first, followed by inverse folding to derive a compatible sequence, and subsequent validation steps, such as refolding to assess structural self-consistency. Despite recent progress toward end-to-end all-atom protein generation (Stark et al., 2025; Butcher et al., 2025), inverse folding continues to play a critical role as a complementary model for refining and validating sequence designs.

A central limitation of inverse folding, and protein design more broadly, is the scarcity of experimentally determined protein structures, which remain costly and time-consuming to obtain (Slabinski et al., 2007; Ding et al., 2022). To mitigate this, several methods augment training data with synthetic structures generated *in silico* by folding models (Hsu et al., 2022; Dreyer et al., 2023; Høie et al., 2024). Because sequence data are far more abundant than structural data, this enables large-scale dataset expansion, typically filtered by folding confidence scores. However, reliance on synthetic structures may bias inverse folding models toward algorithmically favorable and less natural sequence distributions, limiting diversity and deviating from true biological variability.

An alternative way to exploit abundant sequence data is to incorporate language models trained on large protein sequence corpora (Lin et al., 2023; Nijkamp et al., 2023; Ferruz et al., 2022). In the antibody and nanobody domain, such data are particularly abundant due to extensive repertoire sequencing efforts (Olsen et al., 2022). Coupling antibody language models with inverse folding has the potential to improve prediction accuracy while preserving features characteristic of natural antibodies, including favorable developability properties such as solubility, low aggregation propensity, and reduced immunogenicity (Raybould et al., 2019).

Here, we propose combining an antibody-specific language model, which captures natural sequence distributions, with an inverse folder that enforces structural constraints, as illustrated in Figure 1. Instead of fine-tuning inverse folding models on antibody structures, we integrate both components at inference time via ensemble sampling, enabling an effective combination of their complementary signals. We evaluate this approach on highly variable CDR regions, assessing its impact on sequence recovery, structural consistency, and sequence naturalness. By leveraging abundant sequence data instead of limited structural information, this strategy can readily benefit from ongoing advances in language modeling.

**Figure 1.**
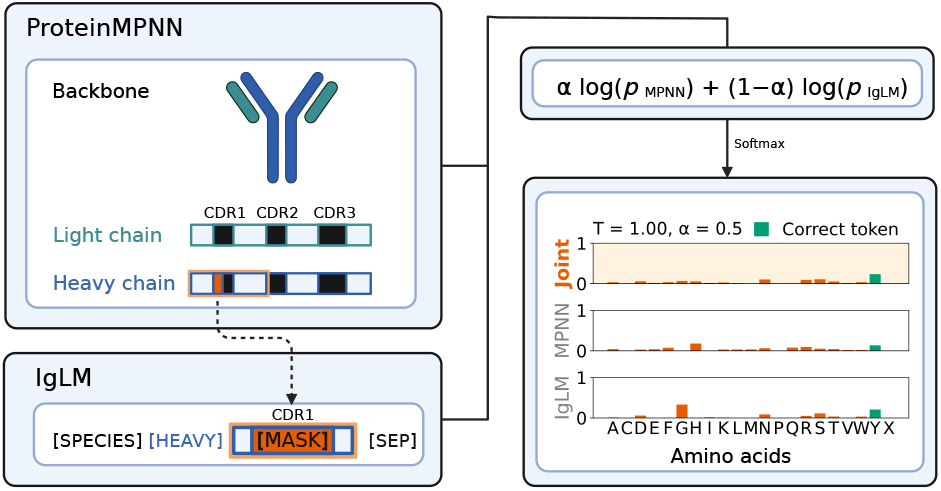
Overview of model ensembling. ProteinMPNN (or AbMPNN) uses the full antibody/nanobody structure, including the target chain and binder framework regions. IgLM predicts one chain at a time, conditioning on the surrounding framework sequence and previously sampled residues for the current CDR. The residue being sampled is highlighted in orange. Log-probabilities from both models (see Methods) are combined and passed through a softmax with temperature T to produce a categorical distribution for sampling the next residue.

## Related Work

### Protein Inverse Folding

leverages diverse machine learning architectures, including graph neural networks (e.g., ProteinMPNN (Dauparas et al., 2022)), sequence-based Transformers (e.g., ESM-IF1 (Hsu et al., 2022)), structure-aware graph Transformer models (e.g., PiFold (Gao et al., 2022)), and discrete diffusion approaches, to predict sequences from structural templates (Ektefaie et al., 2024; Bai et al., 2025). Specialized variants, fine-tuned for properties like solubility (Goverde et al., 2024) or specific protein families, further enhance performance. Antibody-focused inverse folders, including AbMPNN (Dreyer et al., 2023) and AntiFold (Høie et al., 2024), are derived from general protein inverse folders, by continued fine-tuning on antibody datasets. These approaches supplement training with synthetic structures, e.g., from ABodyBuilder2 (Abanades et al., 2023), whereas others, like IgDesign (Shanehsazzadeh et al., 2023), rely exclusively on experimentally determined structures. Discrete diffusion models like AntiDIF (Branson & Deane, 2025) and RADAb (Wang et al., 2024) increase prediction diversity while maintaining high sequence recovery rates.

### Protein Language Models

draw heavily from advances in natural language processing (NLP), with two main paradigms: masked language models (e.g., the ESM family (Lin et al., 2023)) and autoregressive models (e.g., ProtGPT2, ProGen2 (Ferruz et al., 2022; Nijkamp et al., 2023)). These models have been applied to a range of tasks, including sequence annotation, (partial) sequence generation and evaluation, and classification of biological properties. Additionally, they serve as efficient alternatives to time-consuming multiple sequence alignments in protein structure prediction pipelines (Fang et al., 2023). A current trend is the integration of structural and sequence information into multimodal language models, such as ESM3 (Hayes et al., 2025). In the antibody domain, masked language models (e.g., AntiBERTa, AbBERT (Leem et al., 2022; Vashchenko et al., 2022)) are more prevalent and have demonstrated effectiveness in downstream tasks such as paratope prediction (Kalemati et al., 2024). Autoregressive models, such as IgLM (Shuai et al., 2023), generate sequences residue by residue and are designed to produce antibody sequences while assigning likelihood-based scores that reflect their naturalness, making them well-suited for both sampling and sequence scoring.

### Integration of Language Models with Inverse Folding

has already been explored in the past. One prominent approach is LM-Design (Zheng et al., 2023), a framework that combines ProteinMPNN’s structural encoding with a language model (specifically ESM2). This integration requires retraining the language model. While LM-Design is presented as a general framework rather than a specific model, it has been fine-tuned on antibody structure data, as shown by IgDesign (Shanehsazzadeh et al., 2023). Our approach differs from this method in that it requires no retraining, operates purely at inference time, and leverages an antibody-specific rather than a general protein language model. Modern multimodal models such as ESM3 can perform inverse folding without additional training, as they jointly model sequence and structure and are trained on large-scale protein sequence datasets, including antibodies.

## Method

To study ensembling with an antibody language model during inference, we rely on ProteinMPNNN as the inverse folding component. ProteinMPNN is a compact and computationally efficient inverse folding model with strong performance across diverse protein design tasks and broad adoption in design pipelines (Watson et al., 2023; Frank et al., 2024; Pacesa et al., 2025). We also include AbMPNN, a fine-tuned version of ProteinMPNN trained specifically on antibody structures, to assess whether ensembling remains effective when the structure-based model is already antibody-specific.

For the language model component, we choose IgLM, which is specifically trained on antibody sequences. IgLM supports autoregressive and bidirectional sequence generation and aligns well with ProteinMPNN’s sampling strategy. In addition, IgLM’s log-likelihood scores correlate with ProteinMPNN’s amino acid recovery rates (see Supplementary Material S.3), indicating that it can distinguish between high- and low-quality ProteinMPNN predictions and supports its use in ensembling. By modeling natural antibody sequence distributions, IgLM encourages antibody-like predictions.

ProteinMPNN consists of a backbone encoder and a sequence decoder, trained jointly to maximize the likelihood of sequences given their corresponding structures. During inference, the sequence decoder generates residues autoregressively without a fixed positional order, conditioning each prediction on the backbone structure and previously sampled residues. Residues are sampled from a multinomial distribution, with stochasticity controlled by a temperature parameter.

IgLM, which is built on the GPT-2 architecture (Radford et al., 2019), likewise generates amino acid sequences through autoregressive sampling. IgLM captures bidirectional sequence relationships by masking specific regions of the input sequence and predicting the missing residues in the context of the entire sequence, with the predicted residues appended to the input sequence. Nevertheless, it requires a left-to-right sampling process within one masked-out region. We therefore restrict sampling of ProteinMPNN to a left-to-right order within each chain. This does not affect ProteinMPNN’s sampling quality (see Supplementary Material S.3).

To merge the predictions of ProteinMPNN and IgLM, we denote their log-probability outputs at decoding step *t* as

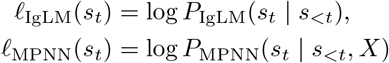

where *s*_*t*_ denotes the amino acid type at step *t*, and *s*_*<t*_ the amino acids sampled at all previous steps, given the left-to-right decoding order. *X* denotes the input structure.

For IgLM, we mask only the CDR under design and provide the flanking framework residues as context.

The ensemble is formed in log-probability space via a convex combination (with coefficient 0 *≤ α ≤* 1):

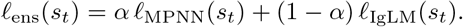

The ensemble distribution is obtained by applying a softmax with temperature *τ >* 0 over all amino acids,

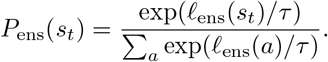

The ensemble can be interpreted as a weighted product-of-experts combination of the model predictions. While the models are not strictly independent due to shared sequence conditioning, this formulation empirically improves sampling performance, as shown in Figure 2 for antibodies and Figure 14 (Supplementary Material) for nanobodies.

**Figure 2.**
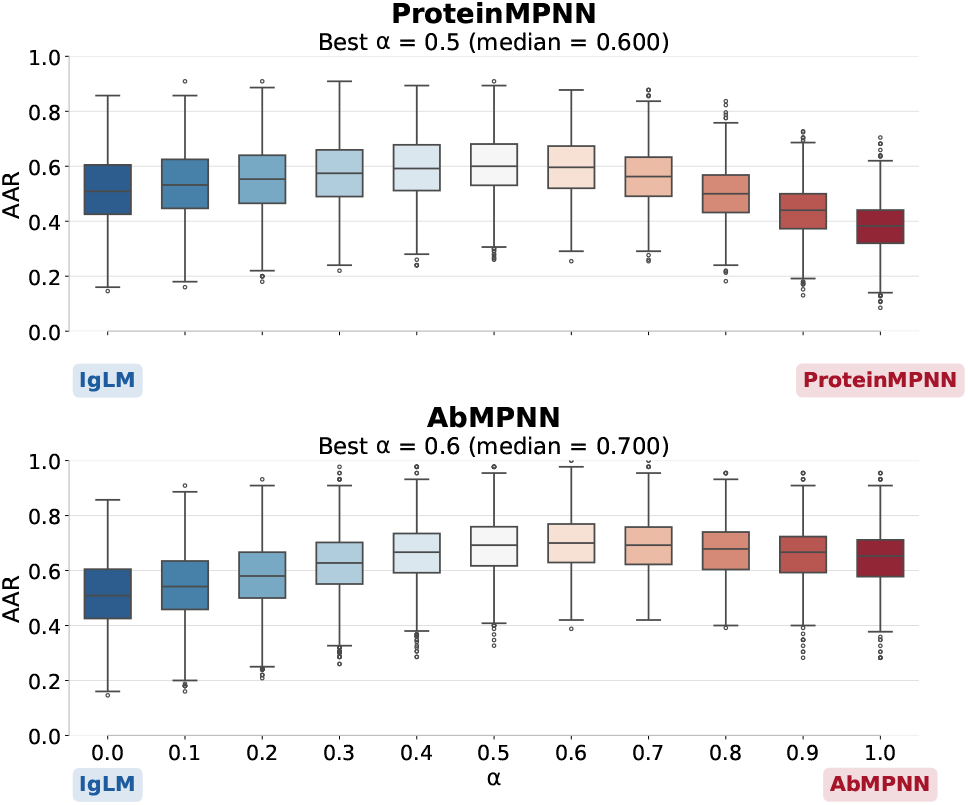
Amino acid recovery rates of CDR loops across different *α* values for ProteinMPNN or AbMPNN ensembles with IgLM, evaluated on antibody validation structures at sampling tempera-ture 0.25 with 10 predictions per structure.

The parameter *α* controls the weighting between the two models, with *α* = 0.5 representing a complete balance between them. We selected *α* empirically based on performance on the validation set.

## Results

We evaluate whether ensembling a general protein inverse folding model (ProteinMPNN) with IgLM can match the performance of antibody-specific models (AbMPNN), and whether AbMPNN also further benefits from ensembling.

ProteinMPNN+IgLM and AbMPNN+IgLM are compared against their base models (ProteinMPNN and AbMPNN) and other inverse folding methods on CDR loop inpainting, targeting the complementarity-determining regions (CDR loops), the most variable regions of antibodies and nanobodies that form the antigen-binding site. Evaluation is performed on a test set of 196 antibody and 93 nanobody structures from the PDB released after ESM3. To reduce redundancy and prevent data leakage, we filter the set such that no pair of structures exceeds 90% sequence similarity with each other, and no structure exceeds 90% sequence similarity to any previously released antibody or nanobody structures available before the cutoff date.

The ensemble weight *α* is optimized using amino acid recovery rate (AAR), a standard sequence-recovery metric in protein design, on a validation set of 1,126 antibody and 374 nanobody structures (Figures 2 and 14 (Supplementary Material)). Tuning *α* for AbMPNN is less straightforward, as it was trained on a substantial fraction of these structures. Nonetheless, ensembling yields consistent but modest improvements in both cases. ProteinMPNN benefits from a smaller *α* than AbMPNN, suggesting that stronger IgLM guidance is advantageous in this setting.

Figure 3a shows that combining ProteinMPNN with IgLM substantially improves CDR recovery, closing the gap to antibody-specific methods such as AntiFold, though still lagging behind AbMPNN, particularly on heavy-chain CDR3. Gains from ensembling AbMPNN with IgLM are smaller but consistent. Nanobody inverse folding remains challenging for all models, reflecting the lower conservation of heavy chains. LM-Design, itself a pLM-based approach, achieves the strongest performance on nanobodies. Overall, ensemble approaches consistently outperform their base models. Figure 3b shows that for ProteinMPNN, ensembling improves AAR while preserving diversity, whereas AbMPNN sees small AAR improvement and increased diversity. Figure 3c reveals a key property. We evaluate predictions from a cumulative-probability perspective, where correctness is defined as whether the ground-truth token appears within the smallest set of most probable tokens whose total probability mass exceeds a threshold p. While AbMPNN achieves high top-1 accuracy, its probability mass is highly concentrated around the top prediction, with negligible mass beyond it. In contrast, the ensembles distribute probability more broadly and more often include the correct token within lower-probability regions, making them more robust for sampling-based design.

**Figure 3.**
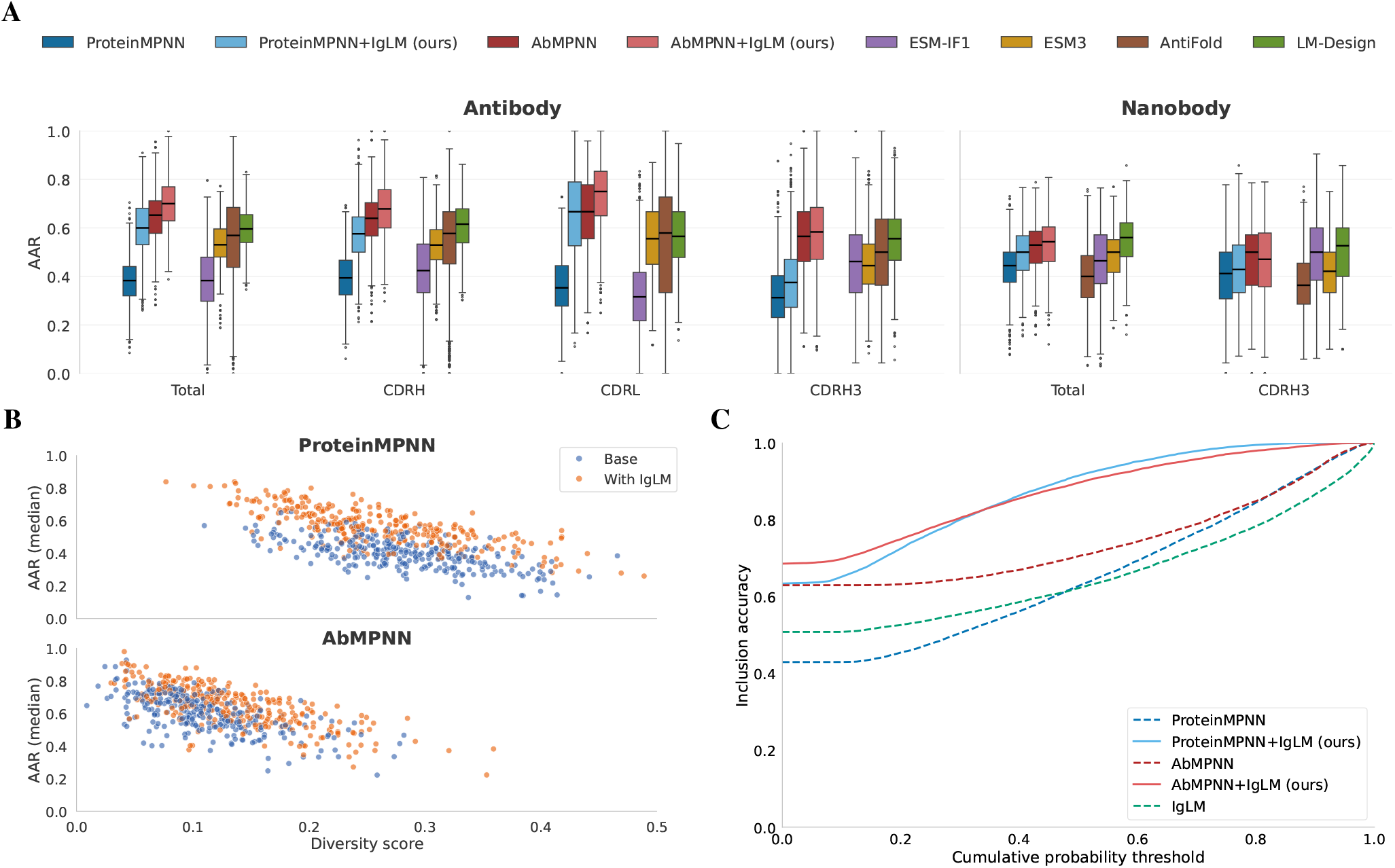
(A) Amino acid recovery rates (AAR) for CDR loops of different models, computed from 10 predictions per sequence on the antibody and nanobody test set at a fixed sampling temperature of 0.25. (B) Diversity score and median AAR per structure, illustrating differences between base and ensemble models for ProteinMPNN and AbMPNN. (C) Fraction of CDR positions where the ground-truth amino acid is included within the set of top-probability tokens whose cumulative probability mass is below threshold p. Curves closer to the upper-left indicate higher confidence in the correct token, as it is recovered at lower cumulative probability thresholds across positions.

Additional analyses in Supplementary Material S.2 show that the ensembles produce highly natural sequences, as measured by ProGen2 and IgLM log-likelihoods, closely matching native sequences, a property not achieved by ProteinMPNN or AbMPNN. We further find that the generated sequences maintain structural consistency, as reflected in Boltz2 (Passaro et al., 2025) refolding experiments, where ensembles consistently rank among the top-performing models in RMSD to native structures. Beyond these metrics, we analyze amino acid profiles and ensembling behavior, focusing on when ensembling is most effective. The ensemble typically follows the more confident model, but reflects agreement when both models are uncertain.

## Discussion

Guiding general protein inverse folders with domain-specific language models improves sequence recovery via inference-time ensembling without modifying model parameters. This approach also maintains sequence diversity and yields more balanced predictions across positions. Structural fine-tuning remains important, as antibody-specific inverse folding models substantially improve challenging regions such as heavy-chain CDR3. Notably, language-model guidance further improves even these specialized models, suggesting the best performance comes from combining structural fine-tuning with complementary sequence-level knowledge from large-scale language models.

### Impact Statement

This work introduces a simple, training-free approach to integrate antibody language models with inverse folding, improving sequence recovery and diversity without requiring additional structural data. By enabling more rapid use of abundant sequence information, this approach may accelerate protein and antibody design workflows, particularly in settings where structural data are limited. Potential applications include therapeutic antibody engineering and biomolecular design. However, as with other generative models in biology, there is a risk that such methods could be misused to design biologically active sequences without sufficient experimental and biosafety evaluation. In addition, model biases inherited from training data may influence generated sequences. Careful wet-lab validation and responsible use remain essential when applying these methods in real-world settings.

## Acknowledgments

This research is funded by the imec.prospect project ADAPT, a research project bringing together academic researchers and industry partners, financed by imec.

