## Supplementary Material for "pLM-Guided Inverse Folding for Antibody Sequence Design"

### S. Supplementary Material

#### S.1. Extended methods

##### Diversity score

We quantified the diversity of our sequence predictions using the metric introduced by (Branson & Deane, 2025) for their discrete diffusion–based inverse folding model.

Sequence diversity $D$ across a set of $M$ predicted sequences $\{\hat{S}_m\}_{m=1}^{M}$ is defined as the average pairwise dissimilarity:

$$D = \frac{1}{M(M-1)} \sum_{j=1}^{M} \sum_{\substack{k=1 \\ k \neq j}}^{M} \Big(1 - \mathrm{SR}(\hat{S}_j, \hat{S}_k)\Big),$$

$$\mathrm{SR}(\hat{S}_j, \hat{S}_k) = \frac{1}{N} \sum_{i=1}^{N} \mathbf{1}\{\hat{S}_j[i] = \hat{S}_k[i]\},$$

where $N$ is the sequence length, and $\mathbf{1}\{\cdot\}$ denotes the indicator function. Here, SR represents the fraction of matching residues between two sequences. This metric captures the average fraction of differing amino acids between predicted sequences and can be applied directly to sequences generated for a given structure.

#### S.2. Analysis of Model Behavior

##### Naturalness

We assess the naturalness of the designed sequences using log-likelihood scores from IgLM and ProGen2 to evaluate whether ensembling with a language model improves sequence quality. IgLM is included as a reference model, as its inclusion in the ensemble is expected to increase log-likelihood values by construction. ProGen2 is included as an independent measure of naturalness, as it has previously been used for this purpose in general protein settings (Gordon et al., 2024).

The analysis shown in Figure 1 demonstrates that, regardless of the underlying base model, ensembling results in designed sequences with log-likelihood scores closer to those of the native sequences. In contrast, sequences generated without ensembling tend to exhibit lower log-likelihoods, indicating reduced naturalness.

##### Structural validation

To evaluate whether sequences designed by the inverse folding models satisfy structural constraints beyond sequence-level agreement (amino acid recovery), we used Boltz2 (Passaro et al., 2025) to refold the designed sequences. Owing to the large number of sequences considered (10 designs per inverse folding model for each PDB structure in the test set, totaling approximately 24,000 sequences), the MSA component of Boltz2 was disabled, and each sequence was refolded only 10 times.

As a reference, we also refolded the native PDB sequences ten times using Boltz2 (Figure 2). The resulting target-aligned binder RMSD values were often relatively high, indicating that Boltz2 does not always reproduce the experimentally observed binding pose. Similar behavior was observed when refolding the native sequences with MSA enabled. While Boltz2 confidence scores were generally consistent with expected ranges and correlated with both target-aligned binder RMSD and framework-aligned CDR RMSD, higher variability was observed for certain structures. Some variability was also observed across repeated Boltz2 runs on the same native sequence, though these differences were limited.

For each structure, we compute two RMSD-based metrics. First, we measure the RMSD between the native binder structure and the designed binder structure after aligning both to the target structure, providing an overall measure of structural agreement in the binding context. Second, we compute a CDR-loop RMSD, where we align the framework regions of the binder and then measure the RMSD over the CDR loops only. This second metric isolates variability in the most functionally relevant regions of the binder while controlling for differences in the conserved scaffold.

While the results, shown in Figure 2, show a small difference favoring the ensemble approach and the base inverse folding models (AbMPNN and ProteinMPNN), the high variance prevents strong conclusions. Differences between nanobody and antibody RMSD values are largely attributable to their differing residue counts and molecular sizes.

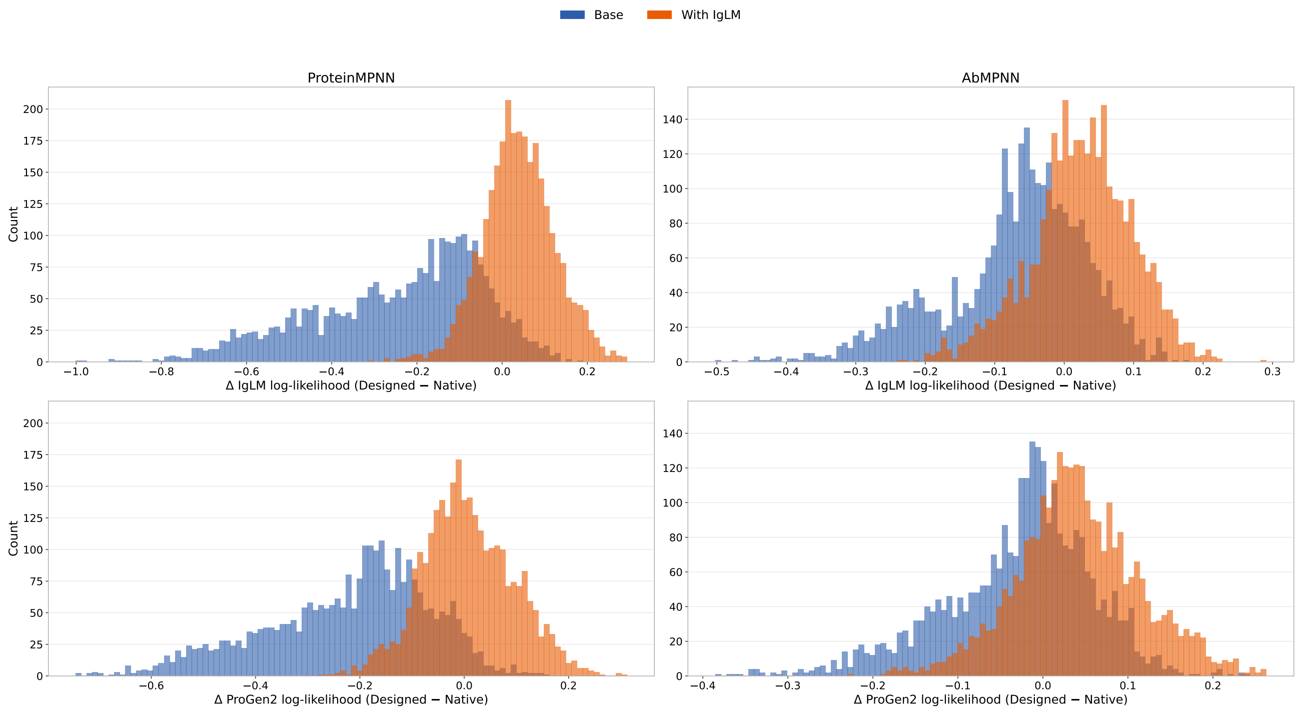


*Figure 1.* Shown are histograms of log-likelihood differences between designed and native sequences for the base models AbMPNN and ProteinMPNN, both with and without IgLM ensembling, based on the antibody and nanobody test set. Log-likelihoods are obtained using IgLM and ProGen2.

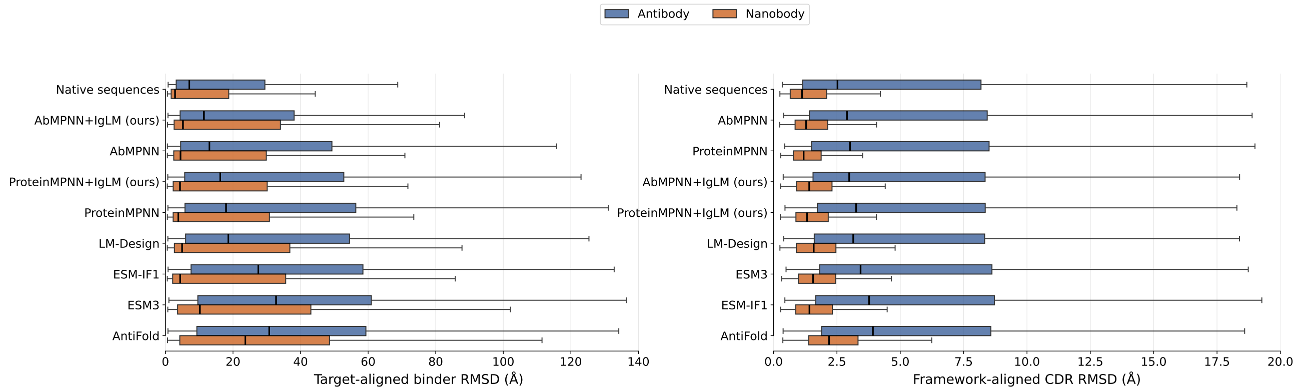


*Figure 2.* RMSD values between 10 Boltz2 predictions for each native and designed sequence (labeled by model name in the plot) in the test set and their corresponding native structures. Target-aligned RMSD is shown for the full binder, while framework-aligned RMSD is shown for the CDR regions.

##### ENSEMBLE MECHANISM ANALYSIS

To better understand when and how ensembling is beneficial, we analyze in more detail the ProteinMPNN+IgLM ensemble, where the effects of ensembling are most pronounced. To do so, we evaluate predictions from ProteinMPNN, IgLM, ProteinMPNN+IgLM, AbMPNN, and AbMPNN+IgLM on the test set under a conditional setting where the subsequence up to the position being predicted is fixed to the correct (native) sequence. This differs from standard autoregressive sampling, where the true (ground-truth) tokens are not available during generation, and sequences are generated solely from previously predicted tokens. This evaluation protocol ensures that all models are evaluated under identical ground-truth context conditioning.

Figure 3 shows the difference in cross-entropy loss between the combined model (ProteinMPNN+IgLM) and ProteinMPNN alone. At temperature T=1 (standard sampling temperature), improvements of ensembling are mainly observed in cases where ProteinMPNN and IgLM strongly disagree, suggesting that the IgLM signal is more informative in these regions. For the lower temperature setting (T=0.25), a different pattern emerges. When the two models strongly disagree (high divergence), ensembling typically improves performance. In contrast, when ProteinMPNN is highly confident (i.e., exhibits low Shannon entropy), its predictions tend to dominate the ensemble, leading to behavior similar to ProteinMPNN alone. Finally, in cases where ProteinMPNN shows high uncertainty but the KL divergence between the two models is low, indicating that IgLM is also uncertain, the ensemble can occasionally degrade performance, as both models provide weak or uninformative signals.

Looking at region-specific signals in Figure 5, the overall trends remain consistent. At T=1, the ensemble cross-entropy loss is often higher than that of the best single model. ProteinMPNN performs better in the CDR3 loops, whereas IgLM achieves lower loss in the remaining loop regions. At T=0.25, however, the ensemble on average outperforms both ProteinMPNN and IgLM across all regions, and is often better than AbMPNN, and comparable to AbMPNN+IgLM.

Overall, these results indicate that the ensemble predominantly combines the strengths of both models by deferring to the more confident prediction. However, there are also cases where ensembling actively improves prediction quality over both models, as shown in Figure 4. In particular, when the correct token is a plausible candidate for both models, but they assign probability mass to different peaks, ensembling can create a new peak in the combined distribution, leading to improved predictions.

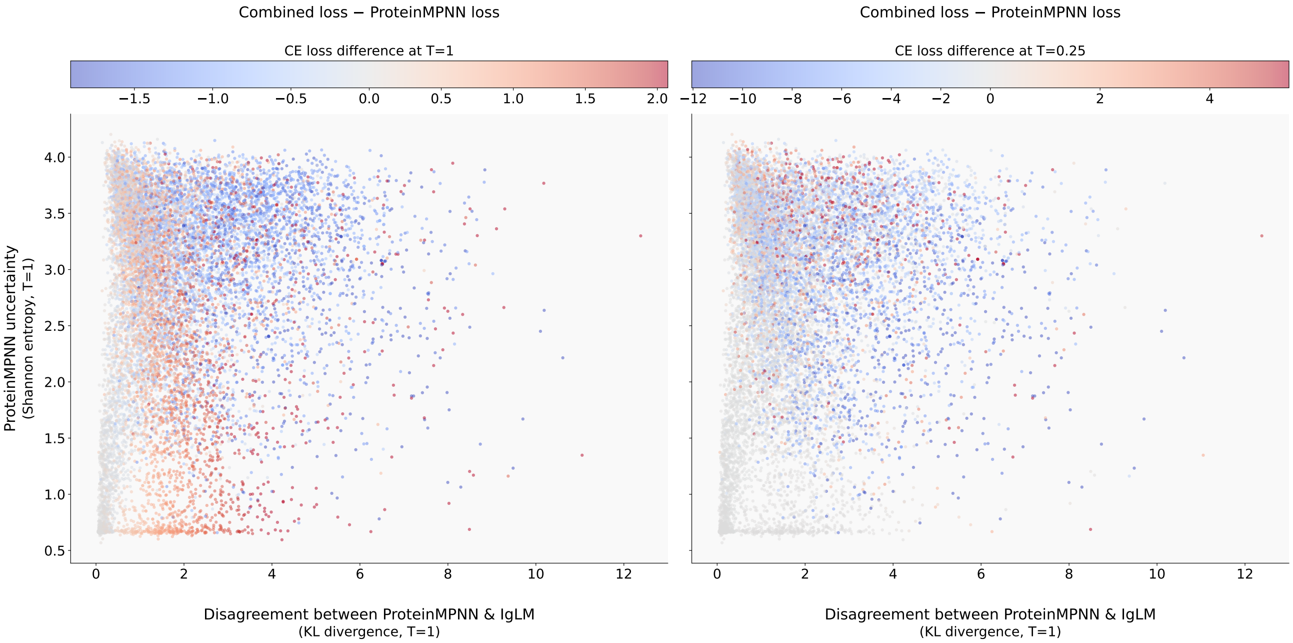


*Figure 3.* Each point in the plot represents the difference in cross-entropy loss between the combined model (ProteinMPNN+IgLM) and ProteinMPNN alone, with blue indicating that the combined model performs better and red indicating that ProteinMPNN performs better. The x-axis shows the KL divergence between ProteinMPNN and IgLM predictions, used as a measure of model divergence, while the y-axis shows the Shannon entropy of ProteinMPNN predictions, used as a measure of model uncertainty. The plot is shown for two different temperature settings (T=1, T=0.25).

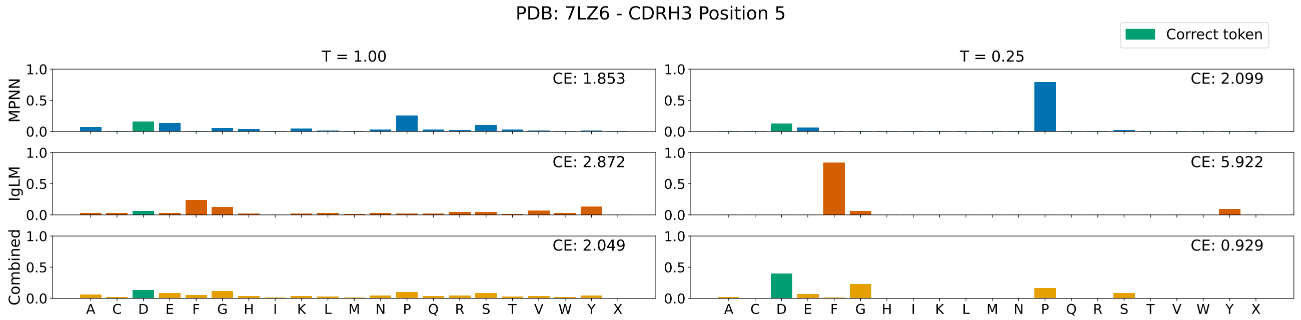


*Figure 4.* Shown is an example of ensembling ProteinMPNN and IgLM, where predictions are first combined at temperature T=1 and then rescaled to T=0.25 by temperature adjustment of the resulting distribution for evaluation. For each model, the predicted categorical distribution over amino acids at a given position is shown, along with the cross-entropy (CE) loss with respect to the ground-truth amino acid. The correct token is highlighted in green. The ensemble redistributes probability mass between the individual models, which can shift the highest-probability token and change the resulting CE compared to the individual predictions. At T=1, the distributions are more diffuse, and the ensemble does not improve CE over ProteinMPNN alone. In contrast, at T=0.25, the sharper distributions lead to more decisive probability shifts, allowing the ensemble to assign the highest probability to the correct token and thereby reduce CE.

###### Amino Acid Frequencies of Models Compared to Native Sequences

We computed amino acid frequencies in the CDR loops of test set structures and compared them to the frequencies in sequences designed by each model, shown in Figure 6. This comparison provides a coarse measure of how well generated sequences reproduce native amino acid composition in antibody binding regions. Ensembling improves ProteinMPNN predictions in terms of matching the native amino acid frequency distribution in CDR loops, but has no positive effect on AbMPNN. The LM-based models LM-Design and ESM3 show high overall similarity to native sequences, although not particularly trained on antibody sequences only. General protein inverse folding models show the largest deviation from native frequencies. Across the better-performing models, there is a tendency to oversample the most frequent amino acids, potentially reflecting training biases toward amino acid recovery objectives.

###### Comparison of amino acid recovery rates across the test set

Figure 7 shows model performance for all CDR loops in the test set.

###### Diversity comparison of model predictions

Figure 8 compares the sequence diversity across models. In general, general protein inverse folding models, which exhibit lower recovery rates, produce more diverse sequences on average. ESM3 shows comparatively low diversity despite its weaker recovery performance.

##### S.3. Ablations and Design Decisions

###### Relationship between amino acid recovery rate and IgLM log-likelihood

Although the correlation in Figure 9 is weak and sometimes negative, amino acid recovery generally correlates with IgLM log-likelihood for both ProteinMPNN and AbMPNN, more strongly in the light chain. Aggregating both models increases the correlation. This suggests IgLM log-likelihood can serve as an empirical proxy for similarity to the native sequence.

###### Comparison of decoding strategies: Random vs. Left-to-Right

In ProteinMPNN, the decoding order is not fixed but is randomly sampled during inference, consistent with the training procedure. Using a left-to-right decoding order, which is required by the ensemble setup, does not significantly affect sequence recovery or sequence diversity in our experiments, see Figure 10.

###### Determination of $\alpha$ for nanobody structures

Tuning of $\alpha$ on nanobody structures, following the same strategy used for antibodies (Figure 11).

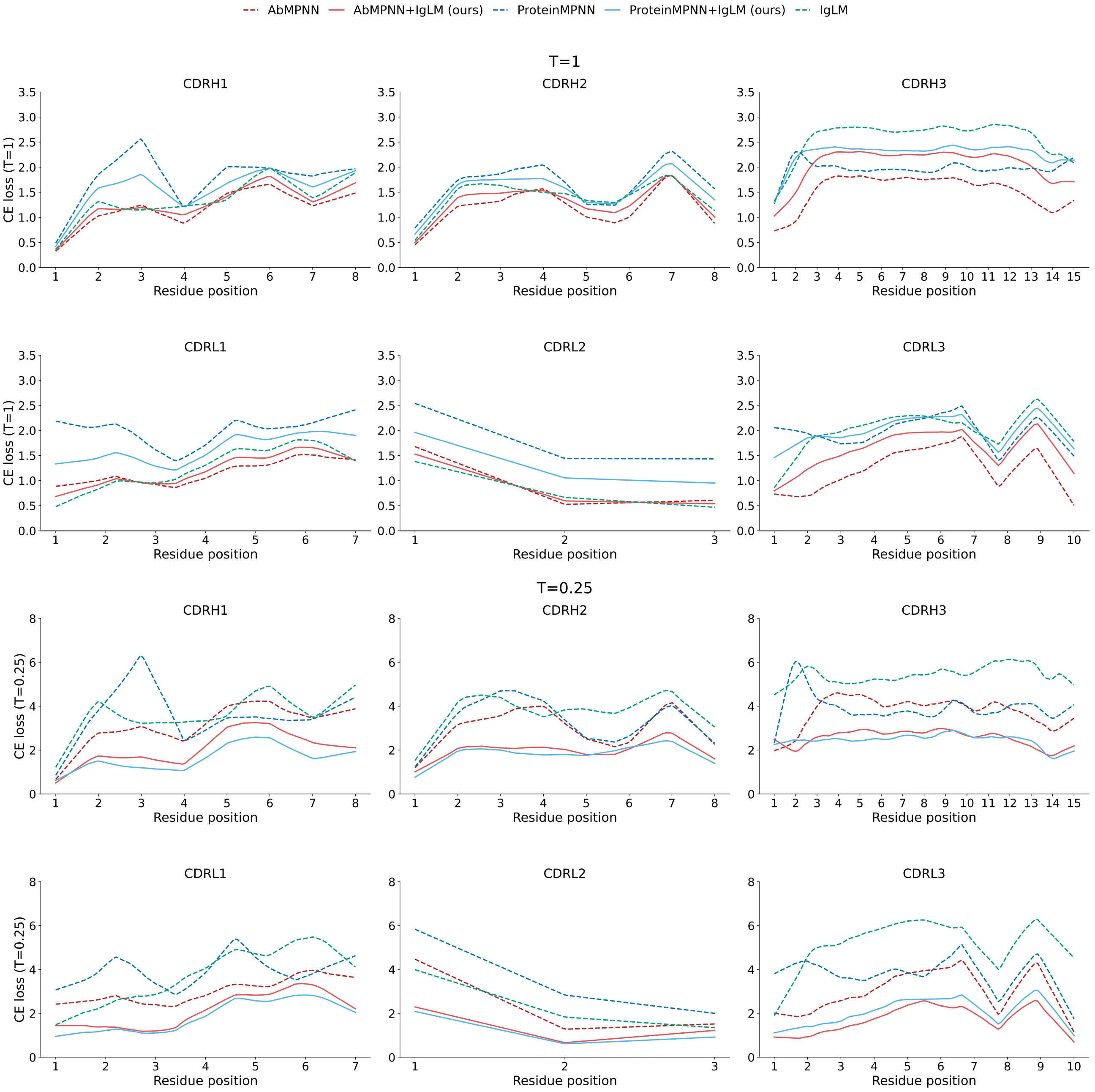


*Figure 5.* The mean cross-entropy loss for different CDR loops is shown for each model at temperatures T=1 and T=0.25. Since CDR loops vary in length, direct position-wise comparison is not possible. To address this, each CDR loop is first normalized to a fixed length by linearly interpolating the cross-entropy loss values onto a grid of 100 evenly spaced positions along the loop. The mean cross-entropy loss is then computed over these 100 normalized positions. Finally, results are rescaled to account for differences in the original average CDR length.

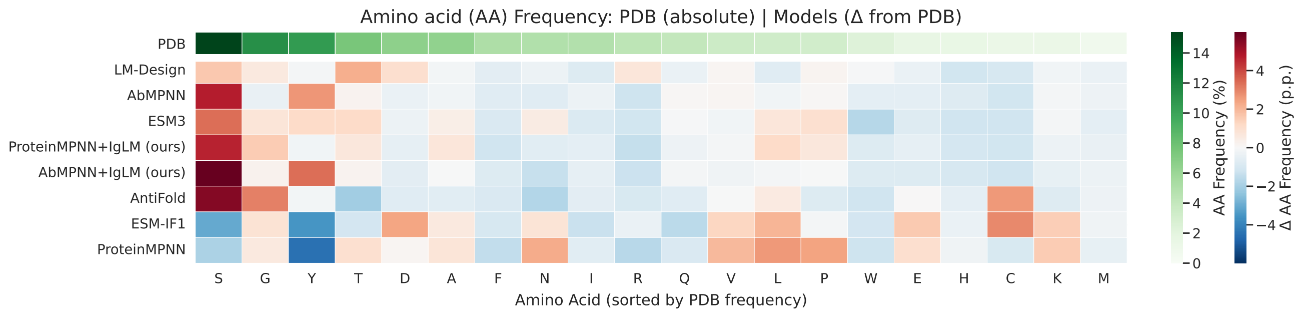


*Figure 6.* Amino acid frequencies in the CDR loops of all test set structures are compared with those predicted by each model. The results are shown as differences relative to the empirical amino acid frequencies observed in the native PDB structures.

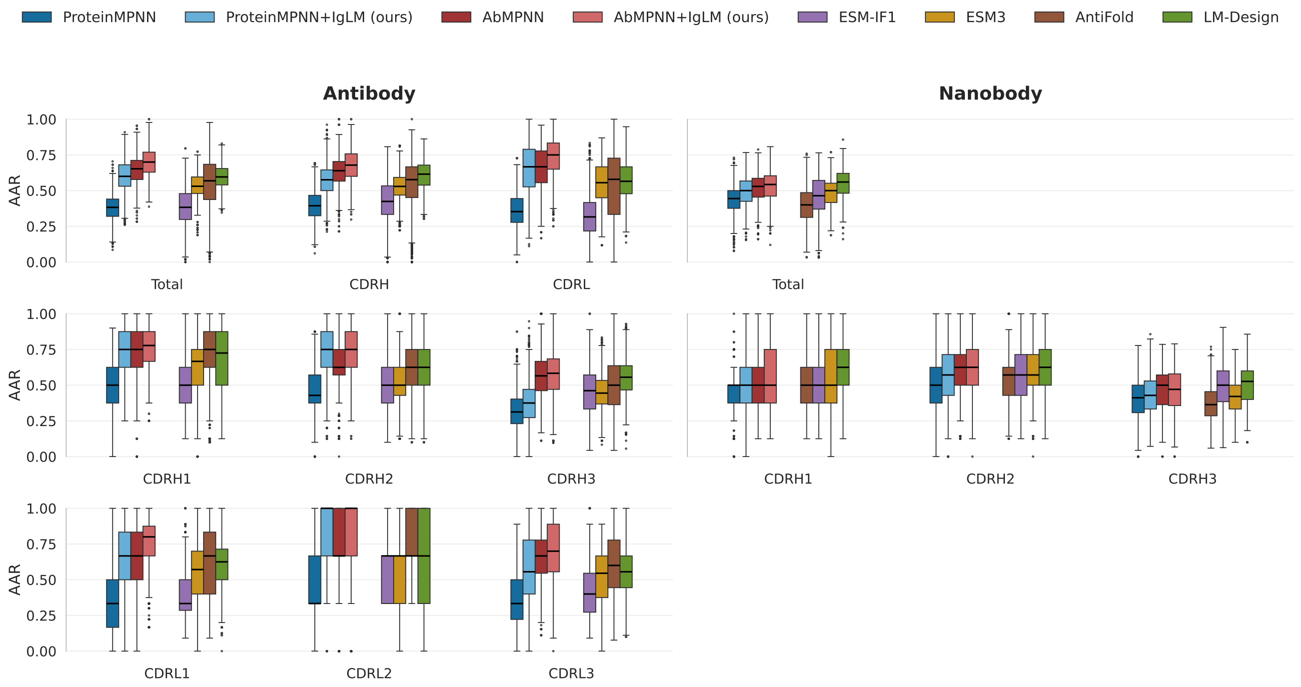


*Figure 7.* Amino acid recovery rates for CDR loops of different models, computed from 10 predictions per sequence on the antibody and nanobody test set at a fixed sampling temperature of 0.25. The plot also reports results separately for each CDR region.

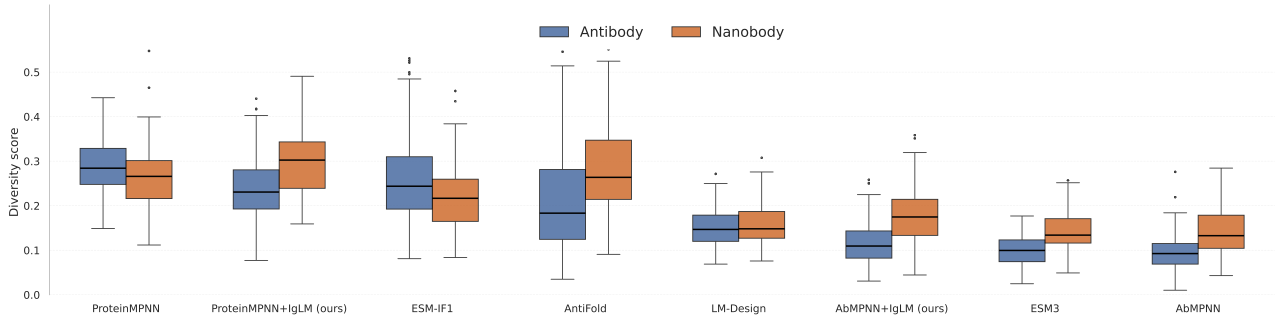


*Figure 8.* Model-wise diversity scores, computed according to Branson et al., for the antibody and nanobody test sets. For each structure, all 10 designed sequences generated by a given model were considered, and a single diversity score was calculated based on the variability among those sequences.

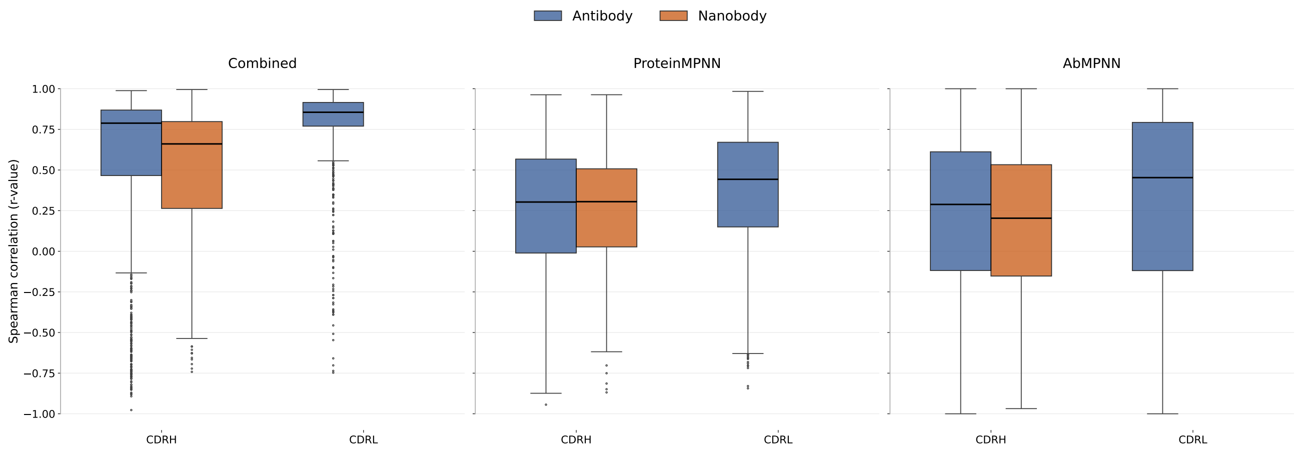


*Figure 9.* Correlation analysis of amino acid recovery rates for AbMPNN and ProteinMPNN predictions on heavy- and light-chain CDR loops, and IgLM log-likelihoods. Results are based on 10 predictions per structure for the nanobody and antibody validation set. Spearman correlations were computed separately for each PDB structure. Boxplots show the distribution for the r-value of correlation analyses.

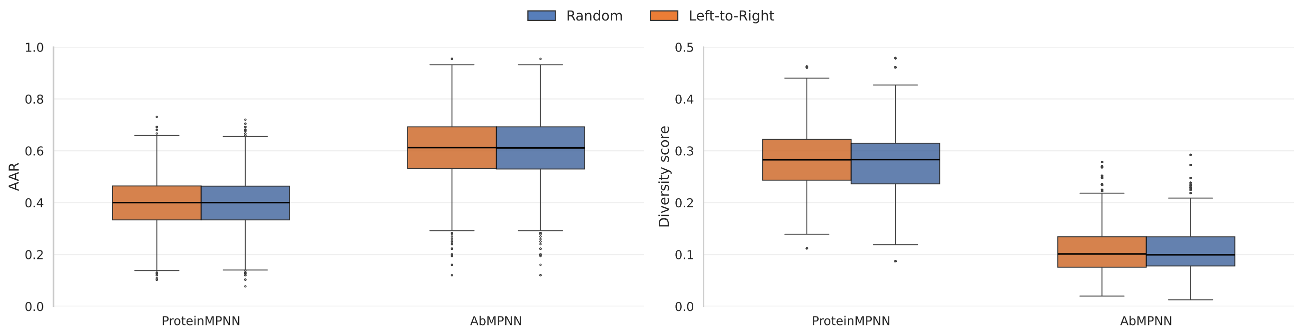


*Figure 10.* Amino acid recovery rates and diversity score for CDR loops under two decoding strategies, random and left-to-right, computed from 10 predictions of ProteinMPNN and AbMPNN on antibody and nanobody test set at a fixed temperature of 0.25.

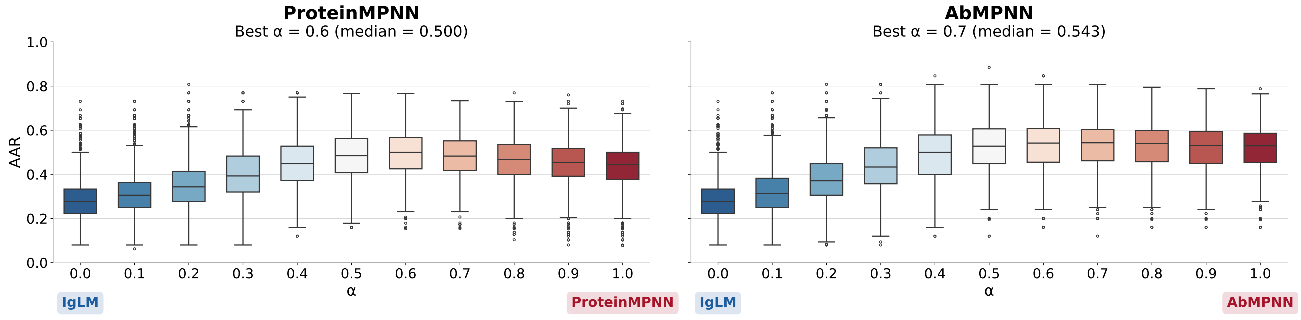


*Figure 11.* Amino acid recovery rates of CDR loops at different $\alpha$ values for ensembles of ProteinMPNN or AbMPNN combined with IgLM, evaluated on nanobody structures from the validation set, with the sampling temperature fixed at 0.25 and 10 predictions per structure.
